# Fly seizure EEG: field potential activity in the *Drosophila* brain

**DOI:** 10.1101/2021.05.10.443444

**Authors:** Atulya Iyengar, Chun-Fang Wu

## Abstract

Hypersynchronous neural activity is a characteristic feature of seizures. Although many *Drosophila* mutants of epilepsy-related genes display clear behavioral spasms and motor unit hyperexcitability, field potential measurements of aberrant hypersynchronous activity across brain regions during seizures have yet to be described. Here, we report a straightforward method to observe local field potentials (LFPs) from the *Drosophila* brain to monitor ensemble neural activity during seizures in behaving tethered flies. High frequency stimulation across the brain reliably triggers a stereotypic sequence of electroconvulsive seizure (ECS) spike discharges readily detectable in the dorsal longitudinal muscle (DLM) and coupled with behavioral spasms. During seizure episodes, the LFP signal displayed characteristic large-amplitude oscillations with a stereotypic temporal correlation to DLM flight muscle spiking. ECS-related LFP events were clearly distinct from rest- and flight-associated LFP patterns. We further characterized the LFP activity during different types of seizures originating from genetic and pharmacological manipulations. In the ‘bang-sensitive’ sodium channel mutant *bangsenseless* (*bss*), the LFP pattern was prolonged, and the temporal correlation between LFP oscillations and DLM discharges was altered. Following administration of the pro-convulsant GABA_A_ blocker picrotoxin, we uncovered a qualitatively different LFP activity pattern, which consisted of a slow (1-Hz), repetitive, waveform, closely coupled with DLM bursting and behavioral spasms. Our approach to record brain LFPs presents an initial framework for electrophysiological analysis of the complex brain-wide activity patterns in the large collection of *Drosophila* excitability mutants.

## Introduction

Studies of the fruit fly *Drosophila melanogaster* have provided many invaluable insights into the genetic basis of nervous system function underlying behaviors (for review, see Sokolowski, 2001 [1]). Such neurogenetic analyses have helped to establish direct links from identified genetic mutations to alterations in nervous system physiology that lead to the behavior of interest. At the behavioral level, a variety of protocols ranging from relatively simple climbing assays (‘negative-geotaxis’) to more sophisticated video analysis of fly behavior have effectively identified and characterized specific abnormalities. In the same mutant flies, alterations in neuronal and synaptic physiology initially uncovered through analysis of peripheral neuromuscular junctions [2, 3, 4, 5], can provide clear readouts on the role of the genes in regulating membrane excitability and synaptic transmission. Recently, a growing number of studies have extended the cellular-level findings, first established in the peripheral nervous system, to by studying central neuronal activity in vivo using patchclamp recording from neuronal soma [6, 7] and imaging with GCaMP or other genetically encoded fluorophores within individual or groups of identifiable neurons [8, 9, 10]. In addition to these robust protocols established to study detailed behavioral and cellular physiological aspects, it is desirable to elucidate global features of ensemble neural activities across various brain regions during behavioral tasks or neurophysiological events.

Pioneered in humans and in other mammals nearly a century ago, electroencephalogram (EEG) techniques provide a readout of different waveforms associated with ensemble and aggregate neuronal activity in different brain regions (Berger, 1929 cited in [11], see also Adrian & Matthews 1934 [12]). EEG signals have found widespread application in revealing key characteristics of global ‘brain states’, such as wakefulness, non-REM and REM sleep [13], and have provided clear hallmarks of epileptiform and other pathophysiological forms of activity [14]. In *Drosophila*, extracellular electrical recordings of activity across neuronal populations have yielded valuable information on sensory physiology, including: photoreceptors in the compound eye (via electroretinogram [15, 16]), olfactory receptor neurons in the antenna (via electroantennogram [17, 18, 19]), and mechanosensory transduction in the Johnston’s organ [20]. A few studies have developed protocols for measuring local field potentials (LFPs) in the brain using glass microelectrodes [21, 22, 23] or multi-electrode arrays [24], focusing on central sensory processing [25, 26], or differentiating between rest, activity and sleep states [27, 28, 29].

In this report, we describe an approach for observing LFP signals from an anatomically specified locus that are suitable for monitoring global electrical activity in the brain during seizures. LFP activity during the stereotypical discharge repertoire of electroconvulsive seizures (ECS) induced by high-frequency stimulation revealed a characteristic pattern with a stereotypic temporal relation to motor-unit discharges at the DLM during behavioral spasms. In contrast, LFP activities during rest or flight could be readily distinguished from seizure-related LFP signals. In the hyperexcitable voltagegated sodium (Na_V_) channel mutant *bangsenseless* (*bss*, [30, 31]), which displays a lower ECS threshold and extreme sensitivity to mechanical shock [32], we found clear temporal alterations in the ECS-associated LFP activity. We further contrasted ECS activity to proconvulsant-evoked spasms by characterizing the LFP activity, demonstrating a unique seizure pattern following injection of the GABA_A_ blocker picrotoxin (PTX). There is a large collection of epileptic *Drosophila* mutants with well-characterized behavioral and electrophysiological phenotypes. Our study provides a first glimpse of a readily accessible brain-wide quantitative signal that can add a new dimension for distinguishing seizure sub-types.

## Methods

### Fly Stocks

The WT strain Canton-S (CS) and bang-sensitive mutant *bangsenseless* (*bss*) were acquired from the Wu Lab’s collection [30]. All flies were reared at room temperature (~22 °C) on Frankel & Brosseau’s cornmeal media ([33], see also [34]).

### Electrophysiology

The tethered fly preparation employed has been described elsewhere [36, 37]. Briefly, flies were immobilized on ice, affixed to a tungsten pin with cyanoacrylate glue (Aaron Alpha Type-203TX), and allowed approximately 30 min to recover. Flies were given a polystyrene ball (~4 mm dia.) to hold and ‘walk’ on during rest periods. Flight activity was induced by gentle air-puffs (see [37] for details).

Flight muscle spikes were monitored by an electrolytically sharpened tungsten electrode inserted into the top-most dorsal longitudinal muscle (DLM) fiber (#45a [38]). A similarly constructed tungsten electrode inserted into the dorsal abdomen served as the reference. Signals were picked up by an AC amplifier (gain: 100x, bandwidth: 1.0 Hz – 10 kHz, AM Systems Model 1800). Electroconvulsive stimulation was delivered following protocols from Lee & Wu [32]. Sharpened tungsten stimulation electrodes were inserted into each cornea (anode in right eye). An isolated pulse stimulator (AM systems Model 2100) generated a 2-s train of 80-V stimuli (0.1-ms duration) at 200 Hz to trigger the ECS discharge repertoire. The stimulation parameters approximately correspond to 1.5x threshold intensity based on previous work [39, 37].

Field potentials were recorded through a low-resistance glass electrode (tip diameter: ~2 μm, resistance: <1MΩ) pulled from filamented borosilicate glass tubing (1 mm OD, 0.58 mm ID, AM systems) with a Brown-Flamming type electrode puller (Sutter P87). These glass electrodes were filled with an adult hemolymph-like saline (108 mM NaCl, 5 mM KCl, 2 mM CaCl_2_, 8.2 mM MgCl_2_, 4 mM NaHCO_3_, 1 mM NaH_2_PO_4_, 10 mM sucrose, 5 mM trehalose, 5 mM HEPES, adjusted to pH 7.5 with NaOH, see [8]). The electrode insertion site was aimed at the center of the three orbital bristles (Figure 1B; see [35]), and the electrode was advanced beyond the cuticle until a stable electrical signal was observed (25-50 μm). During LFP recordings, the abdominal electrode and stimulation electrodes were held at isopotential to each other and served as reference. Field potentials were amplified by a Grass P15 amplifier (gain: 1000x, bandwidth: 0.1 Hz – 10 kHz).

**Figure 1.**
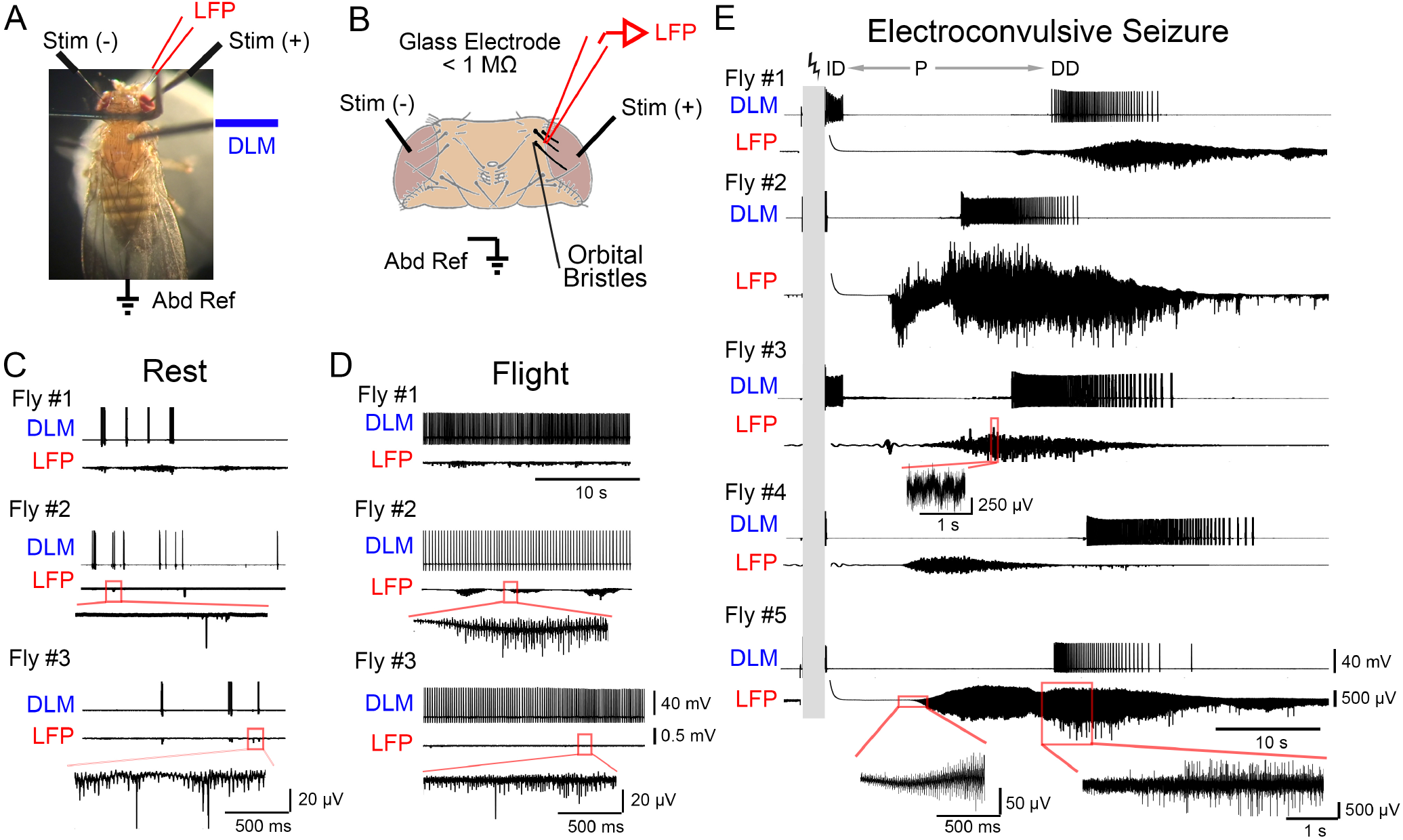
Local field potential measurements in the *Drosophila* brain during rest, flight and electroconvulsive seizures (ECS). (A) Photograph of the tethered fly preparation, with the fly holding a polystyrene ball. The glass microelectrode monitoring local field potential (LFP, red), the tungsten electrode for DLM recording (DLM, blue), isolated stimulation electrodes (Stim (+) and Stim (-), black) across the brain, and abdominal reference electrode (Abd Ref, black) are indicated. (B) Position of the LFP electrode insertion site between the three orbital bristles (bolded in black, drawing adapted from [35]. During LFP recording, both stimulation electrodes are grounded as local current sinks. (C-D) Representative traces of DLM spiking and corresponding LFP activity from three flies resting on a polystyrene ball (C) and flying (D, after dropping the ball). DLM spikes during rest correspond with bouts of grooming activity. Portion of LFP trace from Fly #2 and #3 boxed in red are expanded below the respective traces. (E) DLM spiking and LFP activity during the ECS repertoire. High-frequency, high-intensity stimulation (grey bar, 80 V, 0.1-ms stimuli at 200 Hz for 2 s, see Methods) triggers a stereotypic ECS spike discharge pattern in DLM activity: an initial spike discharge (ID) followed by a paralysis period (P), a delayed spike discharge (DD), and eventual fly recovery. Traces of corresponding LFP activity are shown. For Fly #3 and #5, portions of the LFP trace are expanded below. Note the marked increase in LFP amplitude during ECS, compared to rest and flight.

### Signal Processing and Statistics

For computer-assisted data analysis, DLM spikes and LFP signals were acquired by a USB 6212 data acquisition card (sampling rate: 20 kHz, National Instruments) controlled by a custom-written LabVIEW 2018 script (National Instruments). LFP waveforms were analyzed offline using MATLAB (r2019a) software, including the following procedures. To reduce stimulation-related artifacts, signals were passed through a zero-phase high-pass Butterworth filter (1.0 Hz cutoff, 80 dB attenuation at 0.5 Hz, implemented via the *filtfilt* function). To reduce 60 Hz-related line nose, the signal was then passed through notch filters centered at 60 Hz and the 120 Hz harmonic (± 3 Hz width, 80 dB attenuation). Short-time FFT spectrographs were constructed between 1 and 55 Hz (at 0.5 Hz intervals) using the *spectrogram* function (FFT window width: 1 s, 80% overlap with adjacent windows). Power spectral densities were calculated using the *periodogram* function (0.5 Hz frequency bin width). In Figure 2E, normalized power, *P_N_*, at a particular frequency, *f*, was defined as the fraction of power at a particular frequency within the total power across the 1−55 Hz band, i.e. *P_N_* (*f*) = *P* (*f*)/Σ *P* (*f*). All statistical analyses were conducted in MATLAB.

**Figure 2.**
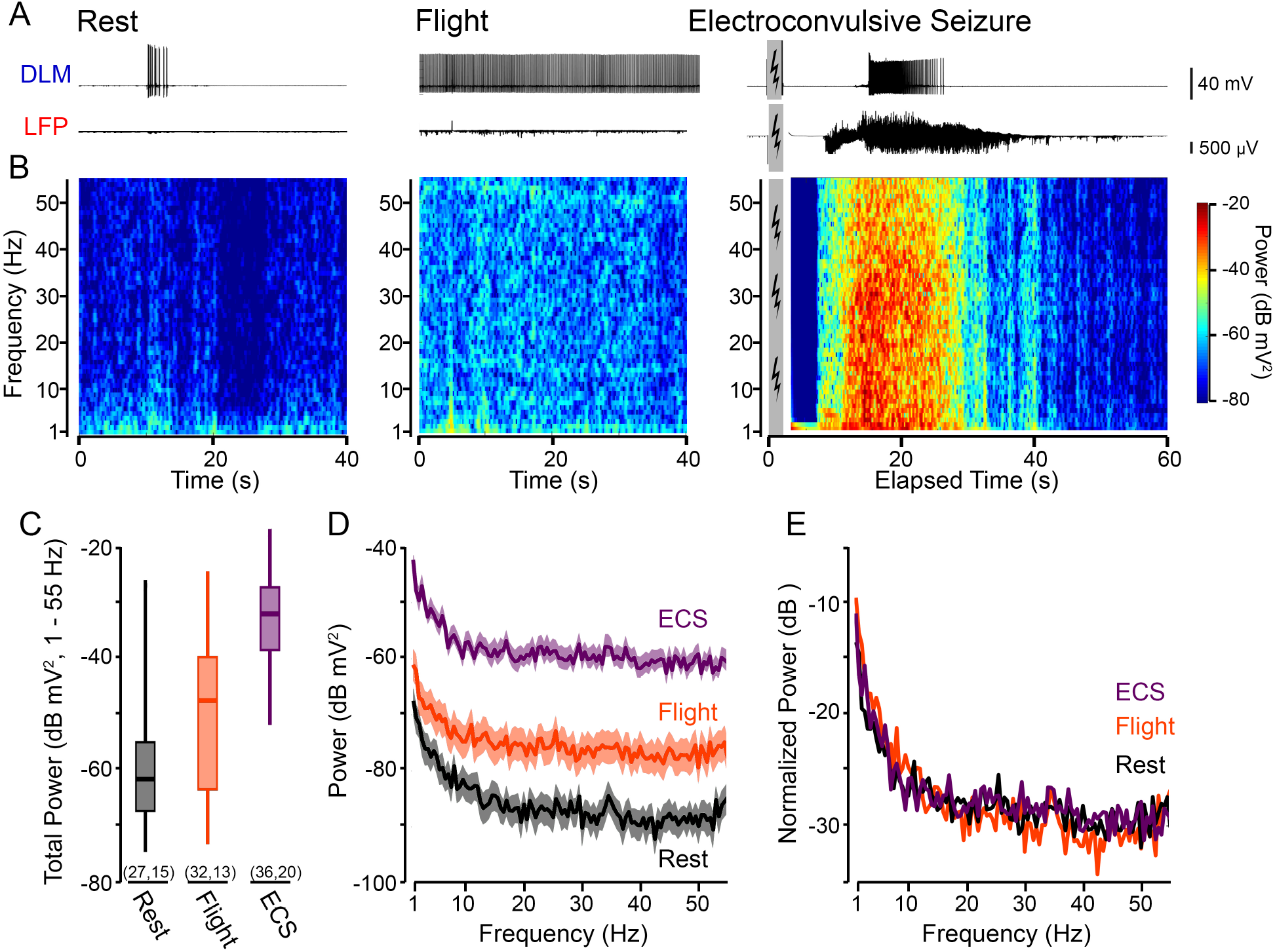
Frequency-domain characteristics of LFP activity related to different behavioral states. (A) Representative traces of DLM spiking (upper traces) and LFP activity (lower traces) during rest, flight and ECS. Grey box denotes the period of electroconvulsive stimulation (ECS trace corresponds to Fly #2 in Figure 1E). (B) Short-time FFT spectrograms corresponding to the LFP activity traces in (A). Window width = 1 s, 80 % overlap between adjacent windows; see Methods for further details. (C) Box plot of the total power in the 1 – 55 Hz range during the respective behavioral states. Boxes represent the 25th, 50th, and 75th percentiles; whiskers the 5th and 95th percentile. Sample sizes in parenthesis indicate the number of trials (left) and flies (right). (D) Averaged FFT spectrogram across all samples of LFP recordings shown in (C) for the respective behavioral states. Solid line indicates the average power across the frequency range; shaded region, SEM. (E) Normalized power, defined as the frequency-specific power divided by the total power (over the 1-55 Hz range), for the respective behavioral states (see Methods).

### Pharmacology

The rapid, systemic drug injection protocol in the tethered fly preparation is described in Lee, Iyengar & Wu (2019, [40]) and was adapted from a dorsal vessel (DV) injection protocol for restrained flies [41]. We utilized a filamented glass electrode similar to the LFP recording electrodes described above, which was filled with solution of the drug of interest dissolved in adult hemolymph-like saline marked with blue #1 dye (16 mg/ml). To inject the solution, the electrode tip was broken and inserted into the DV (see [38]). Positive air pressure, controlled manually through a syringe, pushed the solution (0.33 μL, see [40]) into the DV, and the circulatory system rapidly (~ 10 s) perfused the dyed drug solution.

The nicotinic acetylcholine receptor (nAChR) blocker mecamylamine and GABA_A_ receptor blocker picrotoxin (PTX) were obtained from Sigma Aldrich (Cat#: M9020, P1675 respectively). The voltage-gated sodium channel blocker tetrodotoxin (TTX) was acquired from Cayman Chemical (Cat#: 14964). The concentrations of MEC and TTX were determined empirically to block giant-fiber transmission (1000 and 300 μM respectively). The PTX at the dosage of 100 μM caused a robust sequence spasms described in [40].

## Results

### LFP signals during rest, flight, and electroconvulsive seizure

To record field potentials from the brains of behaving flies, we adopted a tethered fly preparation (Figure 1A, see also [36, 37]) originally developed to record spiking activity in a set of large indirect flight muscles, the dorsal longitudinal muscle (DLMs). During flight, the DLMs undergo isometric stretch-activated contractions at the wing-beat frequency (~200 Hz), with asynchronous spikes (~5 Hz) facilitating Ca^2+^ influx requisite for contraction [42, 43]. Besides flight, several motor patterns drive DLM spiking, including escape [44], grooming [40], courtship [45], and seizure activity [46, 32]. To measure brain field potentials in this preparation, we inserted a stubby, low-resistance (<1 MΩ) glass electrode (“probe electrode”) filled with adult hemolymph-like saline [8] into the head at the recording site (Figure 1A). The three orbital bristles served as landmarks to guide electrode insertion (Figure 1B), and the electrode was advanced through the cuticle until we observed a stable potential in the brain (~25-50 μm). It should be noted that the recorded signal is presumably a complex representation of the electrical activity with different major and minor contributions from various body parts between the probe and reference electrodes (tungsten electrodes inserted into each cornea and into the abdomen). Following conventional nomenclature [47], we refer to the signals picked up in this configuration as local field potentials (LFP). In this preparation, it is straightforward to correlate the LFP with DLM spike patterns associated with different motor programs by using an additional tungsten electrode inserted into the muscle fiber.

We first observed LFP and DLM signals corresponding to two behavioral states: rest and flight. During rest periods, flies held onto or ‘walked’ on a polystyrene ball and periodically groomed themselves. DLM activity during these periods was quite sparse, with brief bouts of spiking closely corresponding with grooming-related wing depression events (c.f. [40]). The corresponding LFP activity was largely consistent with previous reports [21], characterized by signals reaching 2-10 μV amplitude (RMS average, variable across flies presumably reflecting current density and electrode positioning) with superimposed sharp spike-like waves of variable size (Figure 1C). Notably, the LFP signal appeared to loosely correlate with grooming behaviors and related DLM spiking, with transient periods of increased LFP amplitude occasionally observed during periods of no DLM spiking.

A gentle air-puff would initiate a startle response during which the fly dropped the ball and displayed sustained flight (operationally defined to be <30 s [37]), while the DLM spiked rhythmically at ~5 Hz. During flight bouts, the LFP signal was qualitatively distinct from rest period signals, displaying an increased in amplitude (~30 μV RMS average) with more frequent bouts of prolonged spiking activity (Figure 1D). Thus, the LFP signal indeed provides clear indications for distinguishing brain state changes between rest and flight.

High-frequency (200 Hz) electrical stimulation across the brain triggers ECS discharges in flies. As shown in Figure 1E, following a 2-s stimulation train, a distinct sequence of seizure activity consisting of behavioral spasms accompanied by DLM spike discharges are recruited: an initial discharge (ID), followed by a period of paralysis (P), a delayed discharge (DD), and eventual recovery (cf. [32]). Immediately following electroconvulsive stimulation, the LFP remained remarkable quiet for several seconds during the paralysis phase. As the ECS repertoire progressed, before DD onset, a general pattern emerged the LFP signal grew by an order of magnitude or more (to ~200 – 1,500 μV RMS) and these enhanced oscillations continued through the DD period. Following the DD initiation, we often observed prolonged bouts of sharp spiking events in the LFP of millivolt-magnitude (see enlarged traces, Figure 1E). Subsequently, the amplitude of the LFP signal gradually decreased, over several tens of seconds to reach baseline levels. The exact temporal correlation between the LFP oscillations and DD spiking may vary among different experiments, as the ECS-evoked DLM spiking parameters reflect thoracic ganglion driven activity [32] whereas the LFP events are confined to the head. Taken together, these observations provide a robust read-out of central hyper-excitability and hyper-synchronicity which give rise to these large amplitude extracellular field signals that accompany the behavioral and motor unit activity sequence recruited by electroconvulsive stimulation.

### Frequency-domain characteristics of the LFP signal during rest, flight and the ECS discharge repertoire

To examine the frequency components that comprised the LFP signal during the respective motor activities, we applied the Short-Time Fourier Transform (stFFT, over 1-s windows, see Methods) to construct spectrograms across a frequency range (1−55 Hz, representative spectrograms shown in Figure 2A-B). Across the population, we found the total LFP power over the frequency range during ECS discharges was considerably greater than the power during flight or rest (Figure 2C, median values of −32.1, −47.7 and −61.9 dB mV2 respectively). Consistent with previous reports [21], we found the LFP signal during rest displayed increased power in the 1-10 Hz range compared to other frequencies (Figure 2D). Notably, this trend remained true during the flight and ECS discharge activities, in spite of the broad ‘up-shift’ in the power spectrum curve observed. When normalized to total power (Figure 2E, see Methods for computational details), the power spectrum curves during the three motor activities displayed similar inverse-frequency scaling (*P* ∝ 1/ *f^n^*) as reported for brain field potentials from a wide range of species [47]. Linear regression of the log-transformed spectra indicated values of n ranging from 0.72 – 1.05 across the 1 – 55 Hz band width (Supplemental Figure 1).

### Pharmacological manipulation of the LFP signal: effects of blocking Na_V_ channels or cholinergic neurotransmission

The LFP signal we observed was likely a product dominated by brain field potentials, with some other bioelectric phenomena (e.g. heart beats) and system noise picked up between the recording and reference electrode. To identify the origins of LFP signals, we sought to determine how activity patterns were altered by blocking action potential propagation or synaptic transmission. Using a dorsal vessel (DV) drug injection technique [41, 40], we systemically applied the Na_V_ channel blocker tetrodotoxin (TTX), or the nicotinic acetylcholine (ACh) receptor blocker mecamylamine (MEC). Within seconds of injection of TTX, we observed behavioral paralysis coupled with elimination of grooming or flight activity, and the failure of giant-fiber pathway, i.e. brain-stimulation failed to evoke single DLM spikes or ECS discharges. Remarkably, MEC injection led to a comparable effect. Following TTX or MEC administration, the LFP signals were decreased in power to a similar extent (Figure 3A). Both treatments abolished LFP spiking events (Figure 3A), and across the frequency range examined, the signal power was attenuated by ~20 dB (Figure 3B, note that the pre-injection spectra are comparable to the rest-associated spectra in Figure 2D). The results demonstrate that TTX-sensitive Na_V_ channel-driven brain activities generate the predominant component of the LFP signal recorded. Notably, MEC blockade of nAChR also achieved a similar level of LFP attenuation. As in other insects, it is known that ACh is the primary excitatory neuro-trasmitter in the *Drosophila* nervous system [48]. It is also known that synaptic potentials are more effectively picked-up by extracellular field potential recordings than Na^+^ action potentials [49]. Our data provide an independent line of evidence for a major role of cholinergic system in maintaining the basal brain activity as monitored by our LFP protocol.

**Figure 3.**
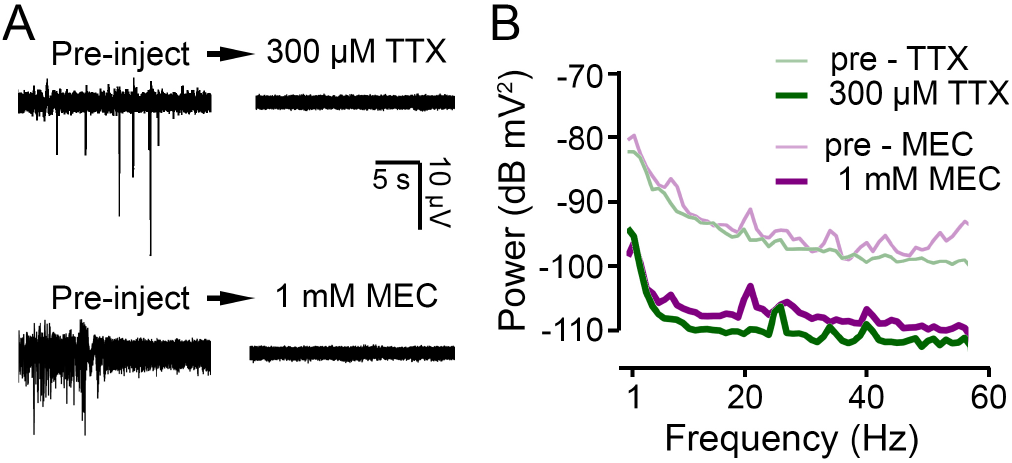
Pharmacological suppression of the LFP signal. (A) LFP traces before and after systemic application of the Na_V_ channel blocker tetrodotoxin (TTX) and the nAChR blocker mecamylamine (MEC) via dorsal vessel injection. Robust effects were observed within seconds, and the traces shown represent the end-point state (achieved ~2 - 4 min). (B) Power spectra of the LFP signals (computed over a 120 s period) before (thin lines) and after injection (thick lines) of MEC (purple) and TTX (green).

### LFP characteristics in *bss*, a ‘bang-sensitive’ mutant

As an initial exploration of the seizure-associated brain dynamics in hyperexcitable *Drosophila* mutants, we examined LFP oscillations during ECS discharges in the ‘bang-sensitive’ mutant allele *bangsenseless* (*bss*, [30]), a gain-of-function allele of the *para* [31], the sole Na_V_ channel gene in *Drosophila* [50]. Upon mechanical shock (e.g. vortexing), *bss* mutants display a striking repertoire of paralysis and vigorous spasms (see [51]). With the ECS protocols, these mutants display a decreased threshold to evoke the DLM spike discharge [46] along with grossly prolonged paralysis interval between the ID and DD [32]. Using the same stimulation protocol for *bss* and the WT counterparts (see Methods), we monitored LFP activity during the ECS discharge sequence. Figure 4A shows LFP oscillations accompanied the DLM spike discharges in *bss* flies. Both *bss* and WT flies displayed similar oscillation frequency characteristics, consisting of a broad increase in power across the 1 – 55 Hz band, but over different timespans (cf. Figure 2B). The episodes of *bss* LFP oscillations were substantially longer than WT individuals (Figure 4C, 87.0 ± 18.4 vs 54.6 ± 5.4 s), corroborating timing differences in the DLM spiking repertoire described previously [32, 39]. Furthermore, we noted a qualitative distinction between *bss* mutants and WT counterparts in that the LFP signal largely waned prior to DD onset in *bss* mutants (Figure 4A), while in most WT individuals, the major LFP power overlapped with DD episodes, suggesting a modification of in the coupling between circuits driving the LFP and DLM spiking.

**Figure 4.**
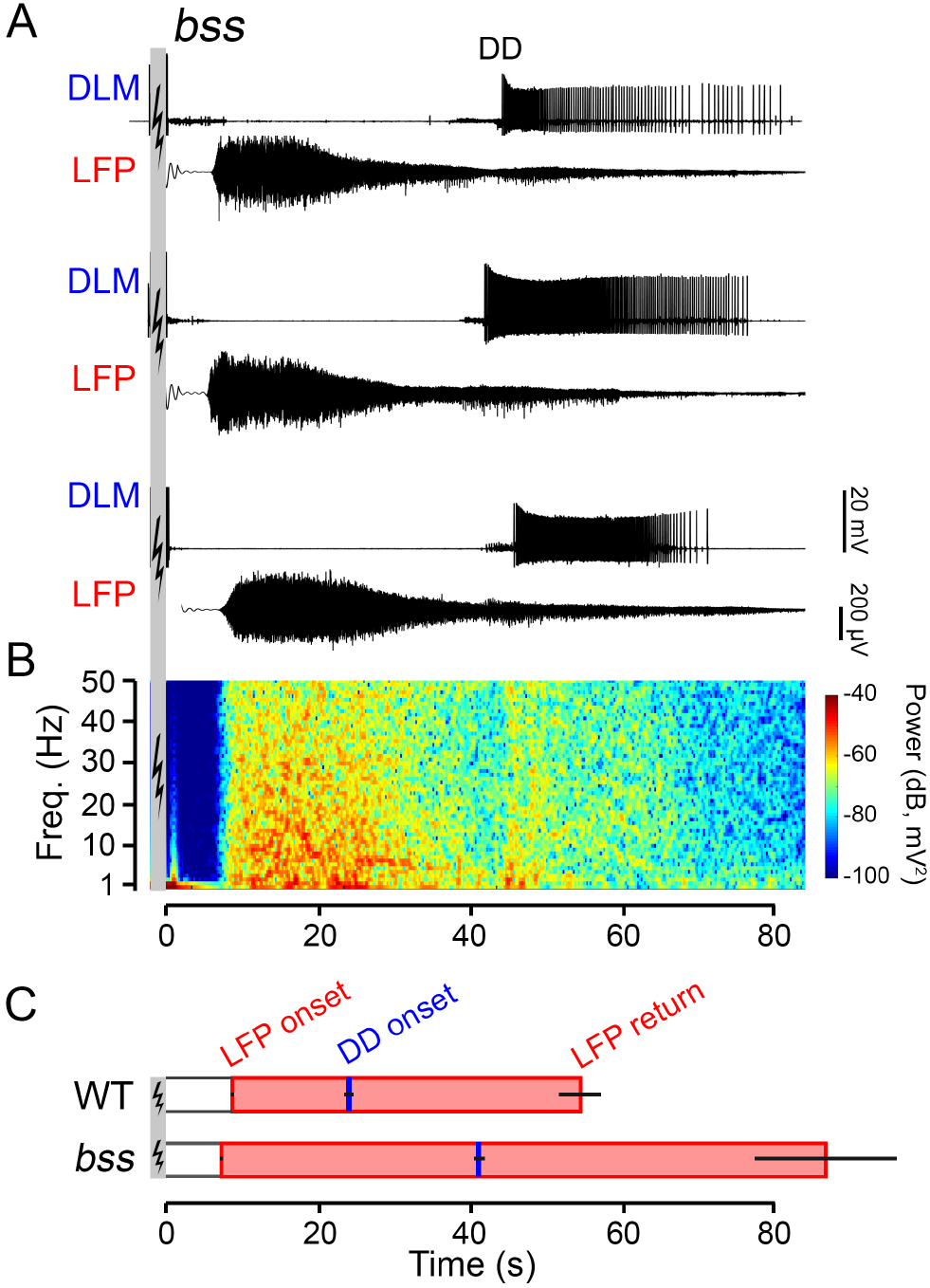
Modifications of the ECS repertoire in *bss*, a hyperexcitable, bang-sensitive mutant. (A) Representative traces of DLM spiking (upper traces) and corresponding LFP activity (lower traces) from three *bss* flies (ECS induction parameters same as Figure 1). (B) Power spectrogram corresponding to the LFP trace directly above in panel (A), constructed in the same manner as Figure 2B. (C) Bar graphs of LFP onset, LFP cessation (shaded pink bar) and DD onset (blue tick) in WT flies compared to *bss* mutants. Note the prolonged LFP signal and delayed DD onset compared to WT flies (cf. Figure 1C). Error bars indicate S.E.M, number of flies as indicated.

### LFP oscillations associated with picrotoxin-induced convulsions

Administration of the GABA_A_ receptor antagonist picrotoxin (PTX) via feeding or injection through the dorsal vessel (DV) triggers a pattern of convulsions and stereotyped DLM spike bursts qualitatively distinct from ECS discharge repertoire [32, 40]. Through monitoring the LFP signal, we identified brain activity patterns induced by PTX. Following DV injection, the LFP signal displayed a sequence of distinct types of oscillations. Within minutes, a stable activity pattern emerged, exhibiting rhythmic depolarization events that coincided with DLM spike bursts and behavioral spasms, which were different from both the rest (pre-injection) state and ECS discharges (Figure 5A vs Figure 2A). Characteristically, LFP activity spectrograms showed strong increases in power around 1 Hz, which corresponded to the DLM burst frequency, while power at higher frequencies remained largely unaltered (representative spectrograms shown in Figure 5B). Thus, the PTX-induced LFP activity reflects a high degree of synchronization correlation between brain activity (LFP waves), motor unit bursting and behavioral spasms. These observations further delineate the proposed distinctions between the modes of seizure activity associated with GABAA blockade and ECS [40].

**Figure 5.**
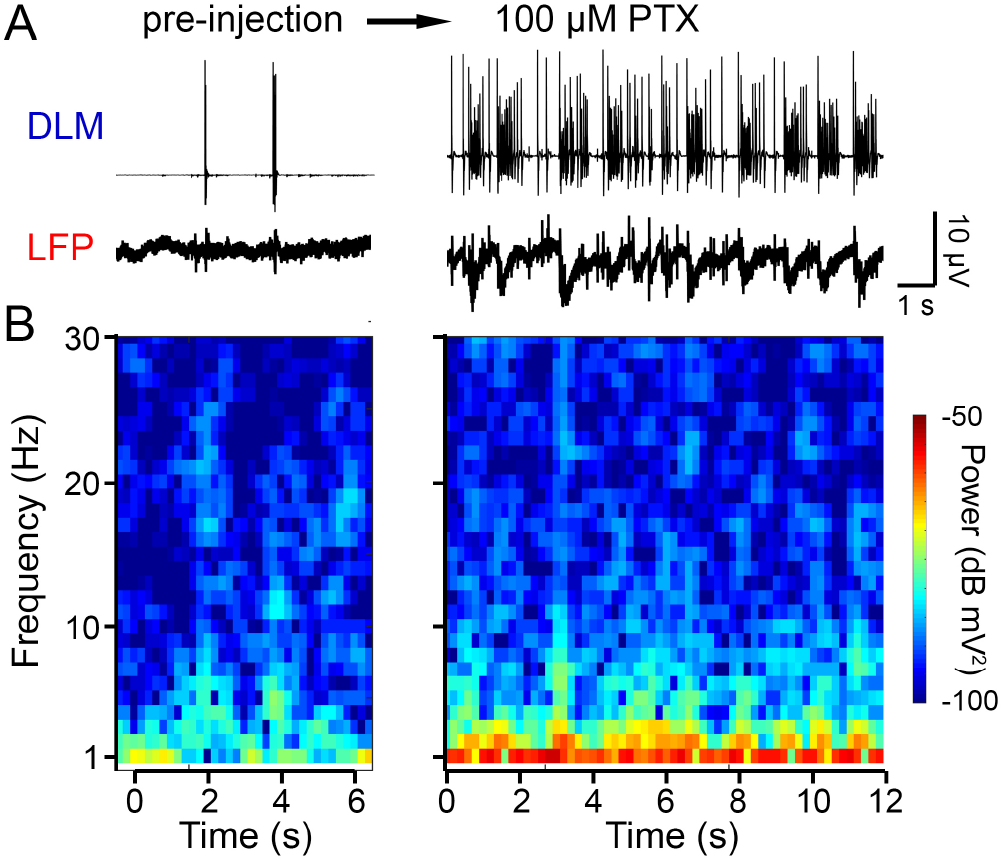
Correlated DLM bursts and LFP waves induced by application of picrotoxin (PTX), a GABAA receptor antagonist. (A) Traces of DLM spiking (upper), and LFP activity (lower) before and after dorsal vessel injection of PTX. The post-injection (~10 min) traces illustrate end-point DLM spike ‘bursting,’ up to ~ 100 Hz within bursts (cf. [40]). (B) Short-time FFT spectrogram of the pre- and post-injection activity. Note the specific increase in low frequency power (about 1 Hz) corresponding to the frequency components of the LFP waves.

## Discussion

Studies of field potential signals from the CNS of *Drosophila* and other insects have contributed to our understanding of olfactory processing [52, 23], attention [25, 53], sleep-wake cycles [28, 54], as well as chill coma and spreading depression events across the nervous system [55, 56]. In this report, we describe a straightforward technique for monitoring event-related LFP signals in the brain during seizure events in tethered behaving flies. Utilizing the orbital bristles [35] as landmarks for electrode insertion, and a defined electrical stimulation-recording configuration (Figure 1A-B), we obtained reliable readouts of global brain signals. Using this approach, we delineated LFP activity during rest from flight-associated patterns. Furthermore, we discovered aberrant yet reproduceable LFP waveforms accompanying the stereotypic ECS discharge (Figure 1E), and proconvulsant (PTX)-induced spasms (Figure 5).

### Origins and properties of the LFP signal: Cellular- and systems-level mechanisms

Field potentials in the brain, arise through the combined current contributions from synaptic transmission and action potential generation as well as other cellular physiological processes [47]. In vertebrate brain slice and *in vivo* preparations, excitatory post-synaptic currents are thought to be the major contributor to LFP signals, as their relatively slow time-course (>10 ms) provides the opportunity for temporal overlap (i.e. synchronization) promoting larger field potential amplitudes [49]. Additionally, currents associated with action potentials and sub-threshold activity in neurons are also recognized to contribute to LFP waveforms to some extent [57, 58]. Thus, a LFP signal (Figure 1) is a complex representation of electrical activity arising between the glass microelectrode and reference electrode(s) in this report.

Our pharmacological experiments indicate that action potential and synaptic transmission blockade similarly suppressed prominent but irregular spike waveforms and power in the *Drosophila* LFP signal (Figure 3). Indeed, previous examination of temperature-sensitive mutants of *para* (encoding Na_V_ channels) and *shibire* (encoding dynamin) found restrictive temperatures similarly blocked components of LFP signal [21]. Although the specific contributions of action potential or synaptic transmission-associated currents in the *Drosophila* LFP remain to be resolved, these results demonstrate that their collective action represents a major component of the LFP signal.

Interestingly, across the three activity states studied, i.e. rest, flight and ECS discharges, the *Drosophila* LFP signal displayed a general inverse relation between frequency and spectral power, *P*∝1/ *f^n^* with *n* ~ 1 (Figure 2D-E). This power scaling phenomenon is also observed across a wide array of vertebrate EEG and LFP from in vivo and brain slice preparations under a variety of physiological conditions, albeit with *n* ~1 - 4 [59, 60, 61]. Although the basis for the inverse power relation remains to be fully understood, the value of *n* is thought to reflect electrical filtering properties of brain tissue [62] and the network organization of the nervous system [47]. The specific mechanisms giving rise to the power scaling relationship in our signal remain to be further investigated, given the differences in circuit organization between *Drosophila* and vertebrate brains.

The LFP signal readily separates brain activity associated with rest, flight and ECS discharges (Figures 1 and 2). At the baseline rest activity pattern, the LFP signal consisted of a ~10 μV (RMS amplitude) signal punctuated with sharp negative deflections (i.e. small spike waveforms, Figure 1C). During flight, the signal displayed a clear increase in RMS amplitude and in the rate of sharp spike waveforms (Figure 1D). Nevertheless, the most striking LFP signal emerged during seizure activity evoked by electroconvulsion (Figure 1E), with a dramatic increase in amplitude superimposed by large amplitude spike waveforms. PTX injection triggered a separate, but clear activity pattern consisting of ‘slow waves’ (ĩ Hz) synchronized with behavioral spasms (Figure 5). Although these LFP activity modes are clear and robustly reproducible, the precise changes in circuit function driving these activity patterns remain to be elucidated. For example, the identified differences between the rest and flight-associated LFP could involve identified octopaminergic [63, 64] or dopaminergic [65] systems in the CNS that are known to modulate flight behavior [66]. Further systematic manipulations of the microelectrode recording site or configurations coupled with transgenic manipulation of activity within specified neural circuits may provide important information regarding the specific neural circuits and/or brain regions in generating the flight- or seizure-specific alterations in LFP signals.

### Monitoring aberrant brain activity through the LFP

In wild-type flies, seizures can be triggered by high-frequency, electroconvulsive, stimulation across the brain, known as ECS discharges [46, 32], or by pro-convulsant (e.g. PTX) application [67, 40], much like seizure induction models in their vertebrate counterparts [68, 69]. Thus far, electrophysiological characterization of seizure activity in *Drosophila* has largely relied on the DLM motor unit as an faithful readout of seizure-associated activity [46, 70, 32, 39, 71]. Following ECS induction, the fly undergoes the highly stereotypic motor sequence consisting of two bouts of DLM spike discharges with intervening paralysis (Figure 1E, see also [32]). Although the DLM is a widely utilized readout, previous studies have shown its action potential bursts are driven by its motor neuron (DLMn) and are temporally correlated with additional thoracic motor units [46]. Furthermore, these DLM discharges overlap with a phase of increased activity that could be picked up within the cervical connective or along the head-abdomen axis [32]. The LFP recordings here provide an initial glimpse of the brain activity patterns accompanying ECS discharges.

Interestingly, during the ECS discharge sequence, the onset of LFP oscillations occurred during the paralysis phase and was distinct from ID or DD onset times (Figure 1E & Figure 4B). This temporal separation suggests distinct physiological processes driving brain and thoracic expression of seizure activity that drive DLM firing. In decapitated flies both ID and DD spike discharges can still be electrically triggered [32]. Thus, while the DLM discharges represent thoracic circuit activity, the LFP signals reflect the physiological state in a different CNS region, i.e. brain-associated seizure activity in *Drosophila*. This ECS-associated LFP pattern consisted of a substantial increase in signal power (across a broad frequency spectrum) and the emergence of repetitive sharp spike waveforms (Figures 1 & 2). Notably, sharp spike events are characteristic of epileptiform activity from mammalian EEG waveforms [72] and from hippocampal and cortical slice LFPs [73, 74]. It is possible that shared physiological mechanisms generate these LFP spike waveforms during ECS in *Drosophila*.

Notably, PTX injection produces a mode of hyperexcitability distinct from the ECS sequence, first inducing flight-like rhythmic (~10 Hz) DLM spiking that gradually evolves into a pattern of regular burst firing (peak frequency ~ 100 Hz), a hyper-active state lasting for hours (this report, and [40]). In contrast to DLM spikes, the PTX-associated LFP signal displayed a relatively slow (~ 1 Hz) negative waves that appeared to be phase-linked with the DLM bursts (Figure 5). Therefore, in this PTX-induced seizure, there appears to be a high degree of synchronization between brain and thoracic activity. Although the similarities and distinctions in physiological mechanisms underlying ECS discharges and PTX-induced seizures remain to be elucidated, both cases provide strong indications for central hyperexcitability and synchronization during seizure activity in *Drosophila*.

Hyperexcitable neural activities and behavioral phenotypes have been associated with a rich and growing collection of *Drosophila* mutants [75, 30, 76, 51]. Certain mutants, such as the gain-of-function para allele *Shudderer* exhibit spontaneous seizures [77]. In other mutants, seizures can be induced by mechanical shock (e.g *bss* [30]; *slamdance* [78, 79]), high temperature (e.g. *seizure* [80, 81]; *down and out*[82]; *zydeco* [83]) or by nutritional deficiencies (e.g. *sugarlethal* [84]). Indeed, many identified mammalian epilepsy-associated genes [85, 86] have homologs in *Drosophila* with mutant alleles displaying clear modifications of circuit-level excitability and/or seizure-like behaviors. There is a growing collection of *Drosophila* transgenic and/or knock-in lines carrying human epilepsy-associated mutations (e.g. *para^GEFS^* [87]; *para^DS^* [88]; *Dube3a* [89]). Although a systematic analysis across seizure-prone *Drosophila* mutants and genetic constructs is beyond the scope of this report, initial glimpse from the bang-sensitive paralytic allele, *bss* (Figure 4) highlights the potential power of the new LFP approach in refined neurogenetic analysis of epileptic categories and various seizure loci in the CNS of *Drosophila*.

## Acknowledgments

We thank Toshi Kitamoto, and members of the Wu Lab, particularly Tristan O’Harrow and Atsushi Ueda, for their helpful discussions over the course of this project. This work was supported by NIH grants (NS111122, AG051513) to CFW, and an Iowa Neuroscience Institute fellowship to AI.

## References

[1] Marla B Sokolowski. Drosophila: genetics meets behaviour. Nature Reviews Genetics, 2(11):879–890, 2001.

[2] Kazuo Ikeda, Seiji Ozawa, and SUSUMU Hagi-Wara. Synaptic transmission reversibly conditioned by single-gene mutation in drosophila melanogaster. Nature, 259(5543):489–491, 1976.

[3] LY Jan and YN Jan. Properties of the larval neuromuscular junction in drosophila melanogaster. The Journal of physiology, 262(1):189–214, 1976.

[4] Obaid Siddiqi and Seymour Benzer. Neurophysiological defects in temperature-sensitive paralytic mutants of drosophila melanogaster. Proceedings of the National Academy of Sciences, 73(9):3253–3257, 1976.

[5] Chun-Fang Wu, Barry Ganetzky, Lily Yeh Jan, and Yuh-Nung Jan. A drosophila mutant with a temperature-sensitive block in nerve conduction. Proceedings of the National Academy of Sciences, 75(8):4047–4051, 1978.

[6] Rachel I Wilson, Glenn C Turner, and Gilles Laurent. Transformation of olfactory representations in the drosophila antennal lobe. Science, 303(5656):366–370, 2004.

[7] Jason W Worrell and Richard B Levine. Characterization of voltage-dependent ca2+ currents in identified drosophila motoneurons in situ. Journal of neurophysiology, 100(2):868–878, 2008.

[8] Jing W Wang, Allan M Wong, Jorge Flores, Leslie B Vosshall, and Richard Axel. Two-photon calcium imaging reveals an odor-evoked map of activity in the fly brain. Cell, 112(2):271–282, 2003.

[9] William C Lemon, Stefan R Pulver, Burkhard Höckendorf, Katie McDole, Kristin Branson, Jeremy Freeman, and Philipp J Keller. Whole-central nervous system functional imaging in larval drosophila. Nature communications, 6(1):1–16, 2015.

[10] Anne K Streit, Yuen Ngan Fan, Laura Masullo, and Richard A Baines. Calcium imaging of neuronal activity in drosophila can identify anticonvulsive compounds. PLoS One, 11(2):e0148461, 2016.

[11] David Millett. Hans berger: From psychic energy to the eeg. Perspectives in biology and medicine, 44(4):522–542, 2001.

[12] Edgar D Adrian and Bryan HC Matthews. The interpretation of potential waves in the cortex. The Journal of physiology, 81(4):440–471, 1934.

[13] MD et al Erik K. St. Louis. Electroencephalography (EEG): An Introductory Text and Atlas of Normal and Abnormal Findings in Adults, Children, and Infants. American Epilepsy Society, 2016.

[14] Soheyl Noachtar and Astrid S Peters. Semiology of epileptic seizures: a critical review. Epilepsy & Behavior, 15(1):2–9, 2009.

[15] Martin Heisenberg. Separation of receptor and lamina potentials in the electroretinogram of normal and mutant drosophila. Journal of Experimental Biology, 55(1):85–100, 1971.

[16] W Pak. Handbook of genetics, chapter Mutations affecting the vision of Drosophila melanogaster. Plenum Publishing, 1975.

[17] Alexander Borst. Identification of different chemoreceptors by electroantennogram-recording. Journal of insect physiology, 30(6):507–510, 1984.

[18] Champakali Ayyub, Jayashree Paranjape, Veronica Rodrigues, and O Siddiqi. Genetics of olfactory behavior in drosophila melanogaster. Journal of neurogenetics, 6(4):243–262, 1990.

[19] Esther Alcorta. Characterization of the electroantennogram in drosophila melanogaster and its use for identifying olfactory capture and transduction mutants. Journal of neurophysiology, 65(3):702–714, 1991.

[20] Daniel F Eberl, Robert W Hardy, and Maurice J Kernan. Genetically similar transduction mechanisms for touch and hearing in drosophila. Journal of Neuroscience, 20(16):5981–5988, 2000.

[21] Douglas A Nitz, Bruno Van Swinderen, Giulio Tononi, and Ralph J Greenspan. Electrophysiological correlates of rest and activity in drosophila melanogaster. Current biology, 12(22):1934–1940, 2002.

[22] Iori Ito, Maxim Bazhenov, Rose Chik-ying Ong, Baranidharan Raman, and Mark Stopfer. Frequency transitions in odor-evoked neural oscillations. Neuron, 64(5):692–706, 2009.

[23] Nobuaki K Tanaka, Kei Ito, and Mark Stopfer. Odor-evoked neural oscillations in drosophila are mediated by widely branching interneurons. Journal of Neuroscience, 29(26):8595–8603, 2009.

[24] Angelique C Paulk, Yanqiong Zhou, Peter Stratton, Li Liu, and Bruno van Swinderen. Multichannel brain recordings in behaving drosophila reveal oscillatory activity and local coherence in response to sensory stimulation and circuit activation. Journal of neurophysiology, 110(7):1703–1721, 2013.

[25] Bruno van Swinderen. Attention-like processes in drosophila require short-term memory genes. Science, 315(5818):1590–1593, 2007.

[26] Angelique C Paulk, Leonie Kirszenblat, Yanqiong Zhou, and Bruno van Swinderen. Closed-loop behavioral control increases coherence in the fly brain. Journal of Neuroscience, 35(28):10304–10315, 2015.

[27] Bruno van Swinderen. A succession of anesthetic endpoints in the drosophila brain. Journal of neurobiology, 66(11):1195–1211, 2006.

[28] Melvyn HW Yap, Martyna J Grabowska, Chelsie Rohrscheib, Rhiannon Jeans, Michael Troup, Angelique C Paulk, Bart Van Alphen, Paul J Shaw, and Bruno Van Swinderen. Oscillatory brain activity in spontaneous and induced sleep stages in flies. Nature communications, 8(1):1–15, 2017.

[29] Bart van Alphen, Evan R Semenza, Melvyn Yap, Bruno van Swinderen, and Ravi Allada. A deep sleep stage in drosophila with a functional role in waste clearance. Science advances, 7(4):eabc2999, 2021.

[30] Barry Ganetzky and Chun-Fang Wu. Indirect suppression involving behavioral mutants with altered nerve excitability in drosophila melanogaster. Genetics, 100(4):597–614, 1982.

[31] Louise Parker, Miguel Padilla, Yuzhe Du, Ke Dong, and Mark A Tanouye. Drosophila as a model for epilepsy: bss is a gain-of-function mutation in the para sodium channel gene that leads to seizures. Genetics, 187(2):523–534, 2011.

[32] Jisue Lee and Chun-Fang Wu. Electroconvulsive seizure behavior in drosophila: analysis of the physiological repertoire underlying a stereotyped action pattern in bang-sensitive mutants. Journal of Neuroscience, 22(24):11065–11079, 2002.

[33] Frankel A. and Brousseau G. Drosophila medium that does not require dried yeast. Drosophila Information Service, 43:184, 1968.

[34] Junko Kasuya, Atulya Iyengar, Hung-Lin Chen, Patrick Lansdon, Chun-Fang Wu, and Toshihiro Kitamoto. Milk-whey diet substantially suppresses seizure-like phenotypes of parashu, a drosophila voltage-gated sodium channel mutant. Journal of neurogenetics, 33(3):164–178, 2019.

[35] G Ferris. The external morphology of the adult.

[36] Jeff E Engel and Chun-Fang Wu. Interactions of membrane excitability mutations affecting potassium and sodium currents in the flight and giant fiber escape systems of drosophila. Journal of Comparative Physiology A, 171(1):93–104, 1992.

[37] Atulya Iyengar and Chun-Fang Wu. Flight and seizure motor patterns in drosophila mutants: simultaneous acoustic and electrophysiological recordings of wing beats and flight muscle activity. Journal of neurogenetics, 28(3-4):316–328, 2014.

[38] A Miller. The internal anatomy and histology of the imago Drosophila melanogaster.

[39] Jisue Lee and Chun-Fang Wu. Genetic modifications of seizure susceptibility and expression by altered excitability in drosophila na+ and k+channel mutants. Journal of neurophysiology, 96(5):2465–2478, 2006.

[40] Jisue Lee, Atulya Iyengar, and Chun-Fang Wu. Distinctions among electroconvulsion-and proconvulsant-induced seizure discharges and native motor patterns during flight and grooming: Quantitative spike pattern analysis in drosophila flight muscles. Journal of neurogenetics, 33(2):125–142, 2019.

[41] Iris C Howlett and Mark A Tanouye. Seizure-sensitivity in drosophila is ameliorated by dorsal vessel injection of the antiepileptic drug valproate. Journal of neurogenetics, 27(4):143–150, 2013.

[42] Michael H Dickinson and Michael S Tu. The function of dipteran flight muscle. Comparative Biochemistry and Physiology Part A: Physiology, 116(3):223–238, 1997.

[43] Shefa Gordon and Michael H Dickinson. Role of calcium in the regulation of mechanical power in insect flight. Proceedings of the National Academy of Sciences, 103(11):4311–4315, 2006.

[44] MARK A Tanouye and ROBERT J Wyman. Motor outputs of giant nerve fiber in drosophila. Journal of neurophysiology, 44(2):405–421, 1980.

[45] Arthur W Ewing. The neuromuscular basis of courtship song in drosophila: the role of the indirect flight muscles. Journal of comparative physiology, 119(3):249–265, 1977.

[46] Paul Pavlidis and Mark A Tanouye. Seizures and failures in the giant fiber pathway of drosophila bang-sensitive paralytic mutants. Journal of Neuroscience, 15(8):5810–5819, 1995.

[47] György Buzsáki, Costas A Anastassiou, and Christof Koch. The origin of extracellular fields and currents—eeg, ecog, lfp and spikes. Nature reviews neuroscience, 13(6):407–420, 2012.

[48] Paul M Salvaterra and Toshihiro Kitamoto. Drosophila cholinergic neurons and processes visualized with gal4/uas–gfp. Gene Expression Patterns, 1(1):73–82, 2001.

[49] F Lopes da Silva and A Van Rotterdam. Biophysical aspects of EEG and Magnetoencephalogram generation.

[50] Kate Loughney, Robert Kreber, and Barry Ganetzky. Molecular analysis of the para locus, a sodium channel gene in drosophila. Cell, 58(6):1143–1154, 1989.

[51] Martin G Burg and Chun-Fang Wu. Mechanical and temperature stressor–induced seizure-and-paralysis behaviors in drosophila bang-sensitive mutants. Journal of neurogenetics, 26(2):189–197, 2012.

[52] Jing W Wang. Odor-induced oscillatory activity in drosophila cns. The Biological Bulletin, 199(2):170–171, 2000.

[53] Martyna J Grabowska, Rhiannon Jeans, James Steeves, and Bruno van Swinderen. Oscillations in the central brain of drosophila are phase locked to attended visual features. Proceedings of the National Academy of Sciences, 117(47):29925–29936, 2020.

[54] Davide Raccuglia, Sheng Huang, Anatoli Ender, M-Marcel Heim, Desiree Laber, Raquel Suárez-Grimalt, Agustin Liotta, Stephan J Sigrist, Jörg RP Geiger, and David Owald. Network-specific synchronization of electrical slow-wave oscillations regulates sleep drive in drosophila. Current Biology, 29(21):3611–3621, 2019.

[55] Kristin E Spong, Esteban C Rodríguez, and R Meldrum Robertson. Spreading depolarization in the brain of drosophila is induced by inhibition of the na+/k+-atpase and mitigated by a decrease in activity of protein kinase g, 2016.

[56] R Meldrum Robertson, Kristin E Spong, and Phinyaphat Srithiphaphirom. Chill coma in the locust, locusta migratoria, is initiated by spreading depolarization in the central nervous system. Scientific reports, 7(1):1–12, 2017.

[57] Supratim Ray and John HR Maunsell. Network rhythms influence the relationship between spike-triggered local field potential and functional connectivity. Journal of Neuroscience, 31(35):12674–12682, 2011.

[58] Mariano A Belluscio, Kenji Mizuseki, Robert Schmidt, Richard Kempter, and György Buzsáki. Cross-frequency phase–phase coupling between theta and gamma oscillations in the hippocampus. Journal of Neuroscience, 32(2):423–435, 2012.

[59] Walter s Pritchard and Dennis w Duke. Measuring chaos in the brain: a tutorial review of nonlinear dynamical eeg analysis. International Journal of Neuroscience, 67(1-4):31–80, 1992.

[60] Kai J Miller, Larry B Sorensen, Jeffrey G Ojemann, and Marcel Den Nijs. Power-law scaling in the brain surface electric potential. PLoS Comput Biol, 5(12):e1000609, 2009.

[61] Joshua Milstein, Florian Mormann, Itzhak Fried, and Christof Koch. Neuronal shot noise and brownian 1/f 2 behavior in the local field potential. PloS one, 4(2):e4338, 2009.

[62] Claude Bédard and Alain Destexhe. Macroscopic models of local field potentials and the apparent 1/f noise in brain activity. Biophysical journal, 96(7):2589–2603, 2009.

[63] Marie P Suver, Akira Mamiya, and Michael H Dickinson. Octopamine neurons mediate flight-induced modulation of visual processing in drosophila. Current Biology, 22(24):2294–2302, 2012.

[64] Floris van Breugel, Marie P Suver, and Michael H Dickinson. Octopaminergic modulation of the visual flight speed regulator of drosophila. Journal of Experimental Biology, 217(10):1737–1744, 2014.

[65] Sufia Sadaf, O Venkateswara Reddy, Sanjay P Sane, and Gaiti Hasan. Neural control of wing coordination in flies. Current Biology, 25(1):80–86, 2015.

[66] Björn Brembs, Frauke Christiansen, Hans Joachim Pflüger, and Carsten Duch. Flight initiation and maintenance deficits in flies with genetically altered biogenic amine levels. Journal of Neuroscience, 27(41):11122–11131, 2007.

[67] Geoff E Stilwell, Sudipta Saraswati, J Troy Littleton, and Scott W Chouinard. Development of a drosophila seizure model for in vivo high-throughput drug screening. European Journal of Neuroscience, 24(8):2211–2222, 2006.

[68] Lowell A Woodbury and Virginia D Davenport. Design and use of a new electroshock seizure apparatus, and analysis of factors altering seizure threshold and pattern. Archives internationales de pharmacodynamie et de therapie, 92(1):97–107, 1952.

[69] Wolfgang Löscher and Dieter Schmidt. Which animal models should be used in the search for new antiepileptic drugs? a proposal based on experimental and clinical considerations. Epilepsy research, 2(3):145–181, 1988.

[70] Daniel Kuebler and Mark A Tanouye. Modifications of seizure susceptibility in drosophila. Journal of Neurophysiology, 83(2):998–1009, 2000.

[71] Salleh N Ehaideb, Atulya Iyengar, Atsushi Ueda, Gary J Iacobucci, Cathryn Cranston, Alexander G Bassuk, David Gubb, Jeffrey D Axelrod, Shermali Gunawardena, Chun-Fang Wu, et al. Prickle modulates microtubule polarity and axonal transport to ameliorate seizures in flies. Proceedings of the National Academy of Sciences, 111(30):11187–11192, 2014.

[72] E Niedermeyer. Abnormal EEG Patterns: Epileptic and Paroxysmal.

[73] Stephen Francis Traynelis and Raymond Dingledine. Potassium-induced spontaneous electrographic seizures in the rat hippocampal slice. Journal of neurophysiology, 59(1):259–276, 1988.

[74] DAVID A Prince and GF Tseng. Epileptogenesis in chronically injured cortex: in vitro studies. Journal of neurophysiology, 69(4):1276–1291, 1993.

[75] Seymour Benzer. From the gene to behavior. Jama, 218(7):1015–1022, 1971.

[76] Juan Song and Mark A Tanouye. From bench to drug: human seizure modeling using drosophila. Progress in neurobiology, 84(2):182–191, 2008.

[77] Garrett A Kaas, Junko Kasuya, Patrick Lansdon, Atsushi Ueda, Atulya Iyengar, Chun-Fang Wu, and Toshihiro Kitamoto. Lithium-responsive seizure-like hyperexcitability is caused by a mutation in the drosophila voltage-gated sodium channel gene paralytic. Eneuro, 3(5), 2016.

[78] HaiGuang Zhang, Jeff Tan, Elaine Reynolds, Daniel Kuebler, Sally Faulhaber, and Mark Tanouye. The drosophila slamdance gene: a mutation in an aminopeptidase can cause seizure, paralysis and neuronal failure. Genetics, 162(3):1283–1299, 2002.

[79] Meghan Horne, Kaitlyn Krebushevski, Amelia Wells, Nahel Tunio, Casey Jarvis, Glen Francisco, Jane Geiss, Andrew Recknagel, and David L Deitcher. julius seizure, a drosophila mutant, defines a neuronal population underlying epileptogenesis. Genetics, 205(3):1261–1269, 2017.

[80] F Rob Jackson, JANE Gitschier, GARY R Strichartz, and LINDA M Hall. Genetic modifications of voltage-sensitive sodium channels in drosophila: gene dosage studies of the seizure locus. Journal of Neuroscience, 5(5):1144–1151, 1985.

[81] Steven A Titus, Jeffrey W Warmke, and Barry Ganetzky. The drosophila erg k+ channel polypeptide is encoded by the seizure locus. Journal of Neuroscience, 17(3):875–881, 1997.

[82] Tim Fergestad, Harinath Sale, Bret Bostwick, Ashleigh Schaffer, Lingling Ho, Gail A Robertson, and Barry Ganetzky. A drosophila behavioral mutant, down and out (dao), is defective in an essential regulator of erg potassium channels. Proceedings of the National Academy of Sciences, 107(12):5617–5621, 2010.

[83] Jan E Melom and J Troy Littleton. Mutation of a nckx eliminates glial microdomain calcium oscillations and enhances seizure susceptibility. Journal of Neuroscience, 33(3):1169–1178, 2013.

[84] Wanhao Chi, Atulya SR Iyengar, Monique Albersen, Marjolein Bosma, Nanda M Verhoeven-Duif, Chun-Fang Wu, and Xiaoxi Zhuang. Pyridox (am) ine 5’-phosphate oxidase deficiency induces seizures in drosophila melanogaster. Human molecular genetics, 28(18):3126–3136, 2019.

[85] Wayne N Frankel. Genetics of complex neurological disease: challenges and opportunities for modeling epilepsy in mice and rats. Trends in genetics, 25(8):361–367, 2009.

[86] Jeffrey Noebels. Pathway-driven discovery of epilepsy genes. Nature neuroscience, 18(3):344, 2015.

[87] Lei Sun, Jeff Gilligan, Cynthia Staber, Ryan J Schutte, Vivian Nguyen, Diane K O’Dowd, and Robert Reenan. A knock-in model of human epilepsy in drosophila reveals a novel cellular mechanism associated with heat-induced seizure. Journal of Neuroscience, 32(41):14145–14155, 2012.

[88] Ryan J Schutte, Soleil S Schutte, Jacqueline Algara, Eden V Barragan, Jeff Gilligan, Cynthia Staber, Yiannis A Savva, Martin A Smith, Robert Reenan, and Diane K O’Dowd. Knock-in model of dravet syndrome reveals a constitutive and conditional reduction in sodium current. Journal of neurophysiology, 112(4):903–912, 2014.

[89] Kevin A Hope, Mark S LeDoux, and Lawrence T Reiter. Glial overexpression of dube3a causes seizures and synaptic impairments in drosophila concomitant with down regulation of the na+/k+ pump atpα. Neurobiology of disease, 108:238–248, 2017.

