## Supplementary figures and images for "Fly seizure EEG: field potential activity in the *Drosophila* brain"

### Supplemental Figure 1

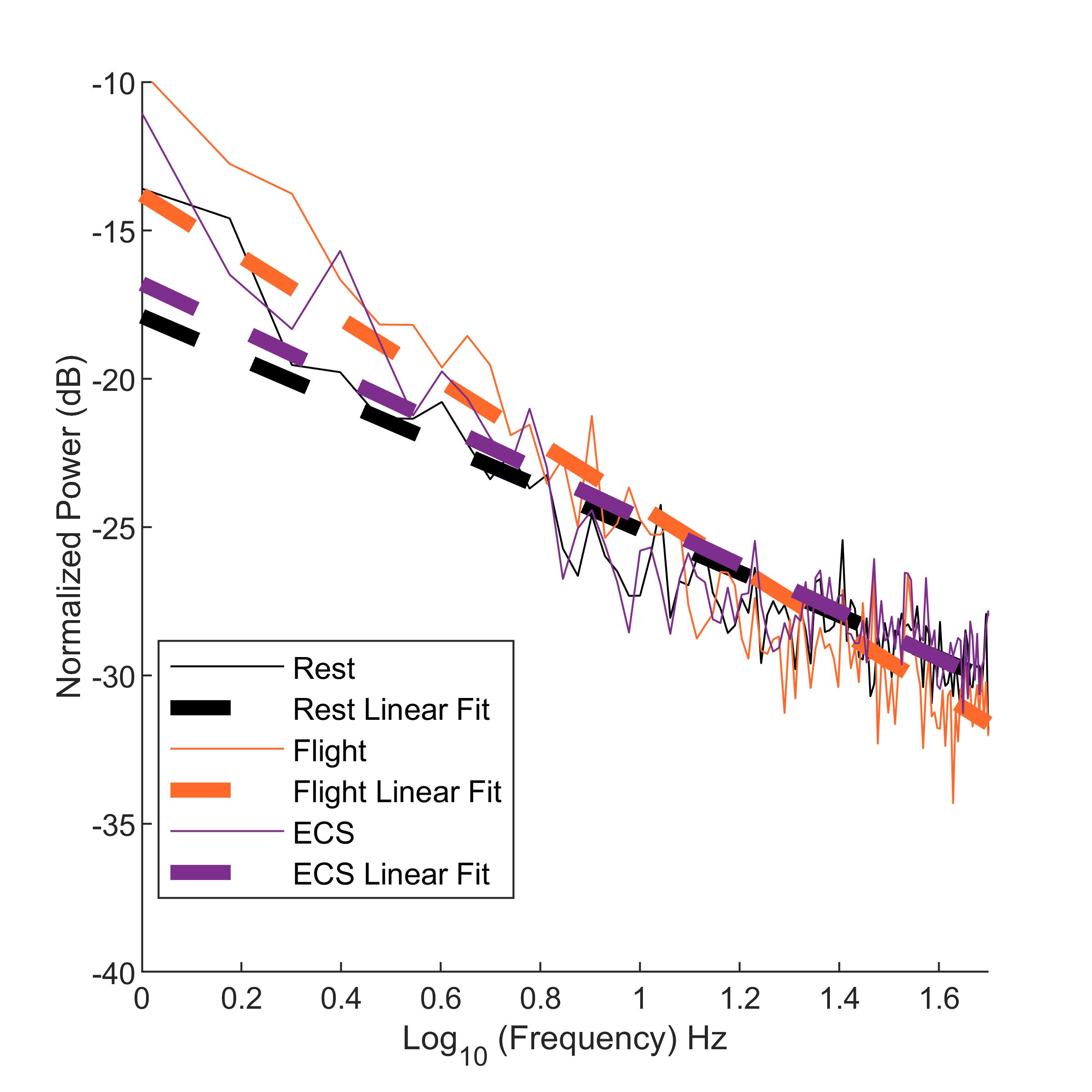
